# Disentangling sources of clock-like mutations in germline and soma

**DOI:** 10.1101/2023.09.07.556720

**Authors:** Natanael Spisak, Marc de Manuel, William Milligan, Guy Sella, Molly Przeworski

**Affiliations:** Department of Biological Sciences, Columbia University, New York, United States; Program for Mathematical Genomics, Columbia University, New York, United States; Department of Systems Biology, Columbia University, New York, United States

## Abstract

The rates of mutations vary across cell types. To identify causes of this variation, mutations are often decomposed into a combination of the single base substitution (SBS) “signatures” observed in germline, soma and tumors, with the idea that each signature corresponds to one or a small number of underlying mutagenic processes. Two such signatures turn out to be ubiquitous across cell types: SBS signature 1, which consists primarily of transitions at methylated CpG sites caused by spontaneous deamination, and the more diffuse SBS signature 5, which is of unknown etiology. In cancers, the number of mutations attributed to these two signatures accumulates linearly with age of diagnosis, and thus the signatures have been termed “clock-like.” To better understand this clocklike behavior, we develop a mathematical model that includes DNA replication errors, unrepaired damage, and damage repaired incorrectly. We show that mutational signatures can exhibit clocklike behavior because cell divisions occur at a constant rate and/or because damage rates remain constant over time, and that these distinct sources can be teased apart by comparing cell lineages that divide at different rates. With this goal in mind, we analyze the rate of accumulation of mutations in multiple cell types, including soma as well as male and female germline. We find no detectable increase in SBS signature 1 mutations in neurons and only a very weak increase in mutations assigned to the female germline, but a significant increase with time in rapidly-dividing cells, suggesting that SBS signature 1 is driven by rounds of DNA replication occurring at a relatively fixed rate. In contrast, SBS signature 5 increases with time in all cell types, including post-mitotic ones, indicating that it accumulates independently of cell divisions; this observation points to errors in DNA repair as the key underlying mechanism. Thus, the two “clock-like” signatures observed across cell types likely have distinct origins, one set by rates of cell division, the other by damage rates.

## I. INTRODUCTION

Mutations are the net result of multiple processes, including endogenous and exogenous DNA damage, errors in DNA replication, and DNA repair. The rate at which they accumulate varies substantially across the human body: mutation rates in germline lineages are typically very low, averaging fewer than one mutation per haploid genome per year, whereas hundreds of mutations accrue yearly in tissues exposed to extensive exogenous mutagens [1–3]. The origins of these pronounced differences remain poorly understood: while the biochemical mechanisms that underlie mutations have been well characterized (e.g., [4, 5]), their relative importance in a given cell type or tissue remains largely unknown.

Over the past decade, a new approach to this question has become feasible, with the analysis of whole genome mutational spectra in cancer genomes and the identification of repeatable patterns of mutations termed mutational signatures [6–8]. In these analyses, single base substitutions (SBS) are classified into 96 types based on the identity of the substitution and the flanking nucleotides, and each tumor sample is characterized by the distribution over the 96 SBS types. Variation across samples is then modeled as a linear combination of relatively few signatures, each characterized by a different profile of the 96 SBS types. The proportion of mutations attributed to a given signature is often described as its “activity” or “exposure” (e.g., [8, 9]), though in what follows, we use the term loading. The idea behind this decomposition is that the signatures reflect one or more mutational processes occurring in different types of samples. In practice, signature profiles and signature loadings across samples are inferred jointly using non-negative matrix factorization [6]. Over 50 signatures have been identified to date [8], which are cataloged and updated in the COSMIC database [10]. Only a subset of COSMIC signatures have a known or partially known etiology, while the sources of most remain elusive. Part of the difficulty is that all signatures reflect the interplay between the initial DNA damage or replication error and DNA repair, and disentangling these from their net effects is not straightforward.

In studies conducted to date, two such COSMIC signatures, SBS signature 1 (SBS1) and SBS signature 5 (SBS5), are ubiquitous. They are detected in tumor samples as well as in healthy somatic cells in humans and other mammals [11], in many cell types contributing the majority of mutations [2]. They are also the predominant contributors to the spectrum of germline mutations [12]. The number of mutations attributed to SBS1 correlates with the age of cancer diagnosis in many cancer types, as do mutations assigned to SBS5, leading the two signatures to be described as “clocklike” [13]. Because they share this behavior, mutations attributed to these two signatures are sometimes combined and analyzed together [14–16].

SBS1 is dominated by cytosine transitions in the CpG context and likely arises from the spontaneous deamination of methylated cytosines [6, 17]. The rate of SBS1 accumulation varies among cell types and increases with the rate of cellular turnover [3, 18]. The loading of SBS1 mutations has also been found to increase in metastatic tumours relative to their primary tumour counterparts, plausibly due to accelerated cell division rates in metastasis [19]. In contrast, SBS5 has a more diffuse distribution among the 96 SBS types. Its ubiquity across cell types, cancerous and healthy, has led to the suggestion that the signature reflects “background” endogenous cellular damage to DNA [20, 21]. However, the number of SBS5 mutations is also affected by the presence of at least one exogenous mutagen, tobacco smoke [22, 23].

Here, we consider conditions under which mutations can accumulate in a clock-like fashion and how the rate of accumulation depends on underlying processes of damage and repair as well as DNA replication. Analyzing whole-genome sequencing datasets of somatic and *de novo* germline mutations in cell types with varying rates of cell division, ranging from post-mitotic neurons to rapidly-dividing intestinal epithelium, we identify mechanisms that likely underlie the clock-like mutation accumulation of SBS1 and SBS5 and highlight key differences between them.

## II. RESULTS

### General model of mutation accumulation

We study the interplay between DNA replication, damage and repair, by extending the model introduced in [24]. We first consider the mutational processes that occur between cell divisions (Figure 1A). A given source of damage causes lesions at a rate *u* per base pair, which are detected and repaired at a rate *r*. Repair leads to mismatches with probability *ϵ*, due to the misincorporation of a nucleotide. We assume that once repair is complete, there are no mechanisms that differentiate the newly synthesized strand from the strand used as a template by the DNA repair polymerase. We further assume that mismatch resolution by mismatch repair or other repair pathways proceeds symmetrically at a total rate *q*, causing a mutation with probability 0.5.

**Figure 1.**
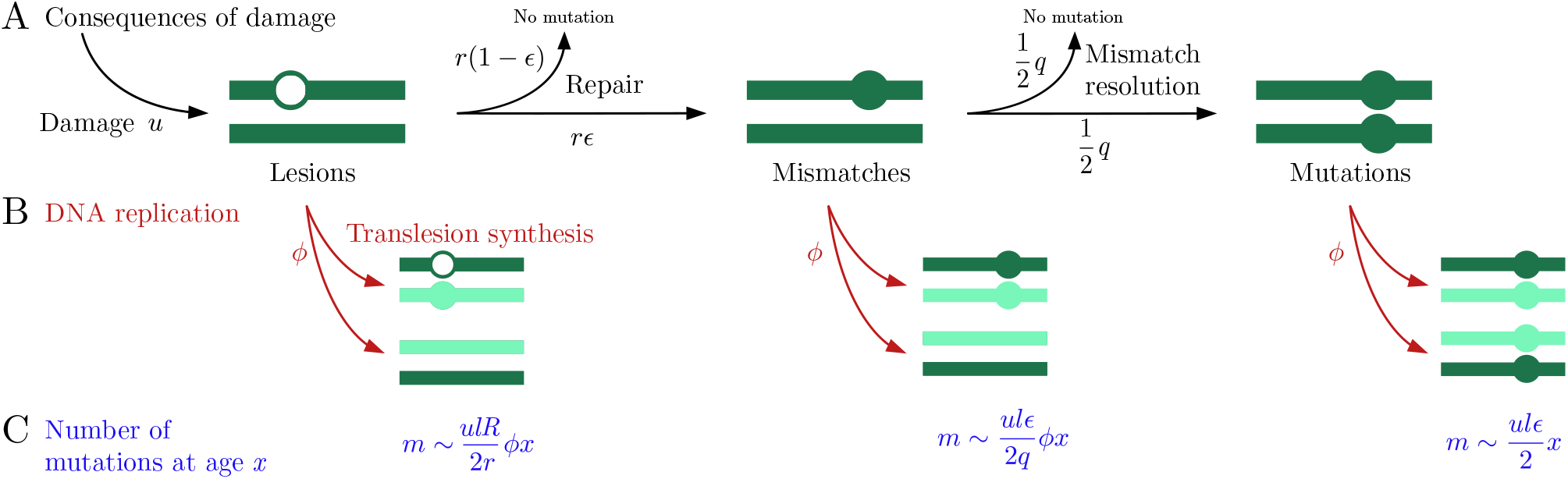
Mutagenic consequences of DNA damage. (A) Interplay of DNA damage and repair during the cell cycle. DNA damage leads to lesions at rate *u* per basepair. Lesions are repaired at rate *r* and lead to mismatches with probability *ϵ*, due to the misincorporation of nucleotides by the DNA polymerase used in repair. Mismatches are resolved symmetrically at rate *q*, resulting in the incorrect basepair and a mutation with probability 0.5. (B) Consequences of DNA replication. Replicating DNA over a lesion requires translesion synthesis. This process is not always accurate, causing an error and a mutation in one of two daughter cells with probability *R* (assuming that the lesion is repaired in the next cell cycle, i.e., that *r* ≫ *ϕ*). Unresolved mismatches cause a mutation in one of the two daughter cells. (C) The predicted number of mutations, *m*, in a genome length *l* at age *x* contributed by the different mechanisms. The genome length is denoted by *l*.

Based on these assumptions, we derive the expected number of lesions, mismatches and mutations and their variances as a function of the time since the last round of DNA replication in a genome of length *l*. The full analysis of the model is presented in Methods; here, we describe the main findings. Over short time periods, *t < r*^−1^, during which repair has had little chance to occur, the number of lesions grows at rate *ul* and the number of mismatches at rate *ulϵ*. Over longer times, *t* ≫*r*^−1^ and ≫ *q*^−1^, the expected number of lesions and mismatches reach a steady state, at which they are approximately equal to *ul/r* and *ulϵ/q*, respectively. In turn, the number of mutations due to incorrect repair increases at rate *ulϵ/*2.

From independent lines of evidence, we know that the vast majority of DNA damage in healthy cells does not lead to mutations: the numbers of mutations in healthy cells are substantially lower than estimated damage rates would suggest [25], and mutation rates in individuals with DNA repair deficiencies are orders of magnitude higher [26]. These observations imply that in healthy cells, the repair rate is on the order of the total damage rate, i.e., that *r* ∼ *ul*. Therefore, the steady state between damage and repair likely is established long before the next cell division. For simplicity, we further assume the rate of mismatch resolution is of the order of the repair rate, *q*∼ *r*. We note that the dynamics of cancer cells may be different from those of the healthy cells on which we focus here; in particular, the repair machinery may be debilitated or overwhelmed and lesions may persist over multiple cell divisions, a phenomenon termed “lesion segregation” [27, 28].

The next part of the model describes the mutational processes that occur during replication (Figure 1B). For simplicity, we assume that the cell divides immediately after DNA replication and treat the two processes as simultaneous, occurring at a fixed rate *ϕ*. An unrepaired lesion stalls DNA replication and triggers the recruitment of polymerases that can replicate over the lesion, through translesion synthesis [29]. This synthesis leads to the incorporation of the incorrect nucleotide opposite the lesion on the template strand with probability *R*, which depends on the type of lesion and possibly on replication timing [30]. If, with probability 1 − *R*, the correct nucleotide is incorporated then, given our assumption that repair is rapid relative to the rate of cell division (i.e., that *r≫ ϕ*), the lesion is likely to be repaired during the next cell cycle. If the translesion polymerase incorporates an incorrect nucleotide, however, the repair process will propagate the error to the complementary strand, generating a mutation. Overall, erroneous translesion synthesis introduces mutations at a rate of *ulRϕ/*2*r*.

Mismatches unresolved during the cell cycle cause a mutation in one of the two daughter cells. Given our assumption that the rate of mismatch resolution far exceeds the rate of cell division (i.e., *q*≫ *ϕ*), these mutations track cell division and accumulate at a rate of *ulϵϕ/*2*q*. In contrast, mutations caused by repair errors are independent of cell divisions and accumulate with absolute time, at a rate of *ulϵ/*2.

Lastly, mismatches also arise during DNA replication due to the misincorporation of nucleotides by replicative polymerases. We denote the probability of a misincorporation per base pair by *w* and the probability that such a misincorporation leads to a mutation by *P* . Importantly, replicating DNA carries transient features that distinguish the newly synthesized strand from its template [31] and help mismatch repair (MMR) substantially decrease the number of replication errors that become mutations, i.e., *P* ≪1. In sum, the number of mutations due to replication errors increases with age at a rate of *wlPϕ*.

Considering these different processes together, our model predicts that the expected number of mutations *m* at age *x* is given by

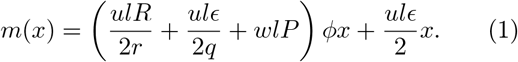

The two terms in this equation correspond to two kinds of clock-like behaviors. The first depends on the cell division rates and includes damage-induced mutations as well as DNA replication errors; assuming, as we do, that cell division rates are fixed, these mutations accrue with time or biological age. The second type of mutation is driven by errors of DNA repair in response to damage; assuming damage rates are constant, it too will depend on absolute time or biological age. However, a distinguishing feature of the two types of clock-like mutations is how they behave as a function of cellular turnover rates; in particular, post-mitotic cells should show no increase of the first type of mutations.

The dynamics of CpG transitions, an important substitution type, can be described by a simpler model, originally introduced in [24]. We consider the consequences of spontaneous deamination of methylated cytosines, which leads directly to a mismatch, and study the kinetics of the reduced model in Appendix A. The expected number of CpG transitions *m* at age *x* is given by

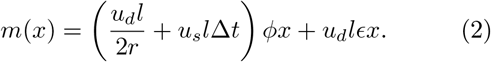

where *u*_*d*_ and *u*_*s*_ denote the deamination rate of methylated cytosines in double stranded and single stranded DNA, respectively, and *l* denotes the number of methylated cytosines. As in the general case (Eq. 1), unrepaired mismatches track the number of cell divisions and errors in mismatch repair depend on absolute time. We also account for the consequences of deamination during DNA replication, which results in mutations tracking cell divisions at a rate proportional to the time period spent single-stranded, denoted by ∆*t*. These results show how a continuous process of deamination may lead to a dependence on cell divisions, as well as a time dependence [24].

In any given cell, a combination of clock-like and nonclock-like mutagenic processes contribute to the mutations. Thus, although in our model, we focus on the conditions under which a clock-like accumulation of mutations will arise, in practice, the total number of mutations of any signature observed in a cell will likely not be strictly proportional to biological age (Eq. 1,2). Instead, there may be a non-negligible contribution of mutations that do not increase with age (i.e., are “age-independent”). These mutations can have several different sources: for example, they may have accrued during the first few cell divisions of development [32], the last stages of cell differentiation [22], or in response to an acute exposure to exogenous mutagens [33]. If these mutations occur in a burst, i.e., over a short period of time, then they will contribute a set number of mutations regardless of age, and a regression of the number of mutations against age can lead to a positive intercept.

A non-zero intercept can also be generated by a clocklike process occurring at a varying pace over development. We discuss model expectations in this case with a toy piece-wise linear model, in which mutations accumulate at two different rates over two time periods (Figure A2): initially, the number of mutations grows at rate *μ*_0_, and after some time *x*_0_, the underlying parameters of mutagenesis (Eq. 1) change and mutations accumulate at rate *μ* ≠ *μ*_0_. For data points collected after time *x*_0_, the expected number of mutations is then

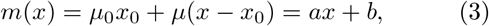

where *a* = *μ* and *b* = (*μ*_0_ −*μ*)*x*_0_. Hence, the intercept is positive if *μ*_0_ *> μ*; for example, if entering a postmitotic state at *x*_0_.

Conversely, an increase in the rate of cell division or the rate of damage at *x > x*_0_ (such as occurs in the lung epithelia of regular smokers [22]) will lead to an increased slope in the number of mutations with age. If the mutation rate was significantly lower at earlier stages (say, before the person smoked), i.e., if *μ*_0_ *< μ*, then a regression of the total number of mutations on age will yield a negative intercept.

In reality, many such effects are superimposed, and it is therefore important, in disentangling the origins of clock-like mutations, to distinguish those that accrue with age from those that contribute to the intercept, and may or may not be clock-like. Concretely, in data sets for which there is a significant positive intercept, the goal is to tease apart signatures that contribute at a constant level in all samples from signatures for which mutations increase in number with age. To this end, we extend the standard signature decomposition method to allow for a mixture of two components, a constant and an age-dependent one, and attribute mutational signatures to the two jointly.

### Clock-like signatures across cell types

In order to gain insight into the origins of mutations that accumulate with age, we analyze patterns of mutation accumulation across cell types with different characteristics. To this end, we consider data sets that provide single-cell resolution mutation data, collected using a variety of experimental approaches (see Methods), including mutations in neurons and muscle cells [3], liver hepatocytes [34], lung epithelium [22], small bowel epithelium [35], colonic epithelium and testis seminiferous tubules [2], as well as germline mutations identified from blood samples of pedigrees [36, 37]. We rely on mutation data from donors without a disease diagnosis and on lung samples from non-smokers (see Methods).

To compare observations with our model predictions, we develop an approach to focus on the subset of mutations in a given cell type that accumulate with age. This is done in two steps. First, we fit a linear model for all mutations jointly, i.e., *y* = *ax* + *b*, where *y* denotes the number of mutations per genome and *x* denotes age (see Figure 2A for mutations in neurons). Second, we model the distribution over the 96 substitution types as a mixture distribution of two components: the constant component (Figure 2B, yellow) contributes on average the same number of mutations in samples of all ages, whereas the age-dependent component (Figure 2B, blue) contributes an increasing number of mutations with age. We decompose the slope and the intercept into COSMIC signatures jointly, such that the dependence of the number of mutations *y*_*s*_ attributed to a given signature *s* on age takes the form

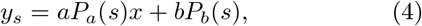

where *P*_*a*_(*s*) and *P*_*b*_(*s*) denote the loadings of signature *s* in age-dependent and constant components, respectively. We estimate the loadings by extending the standard methods of signature attribution (see Methods); see Figure 2C-D for the example of loadings estimated for mutations in neurons. We note that applying this method is only possible if the intercept *b* is large enough (i.e., if the data contains enough age-independent mutations to attribute mutations to signatures in the constant component).

**Figure 2.**
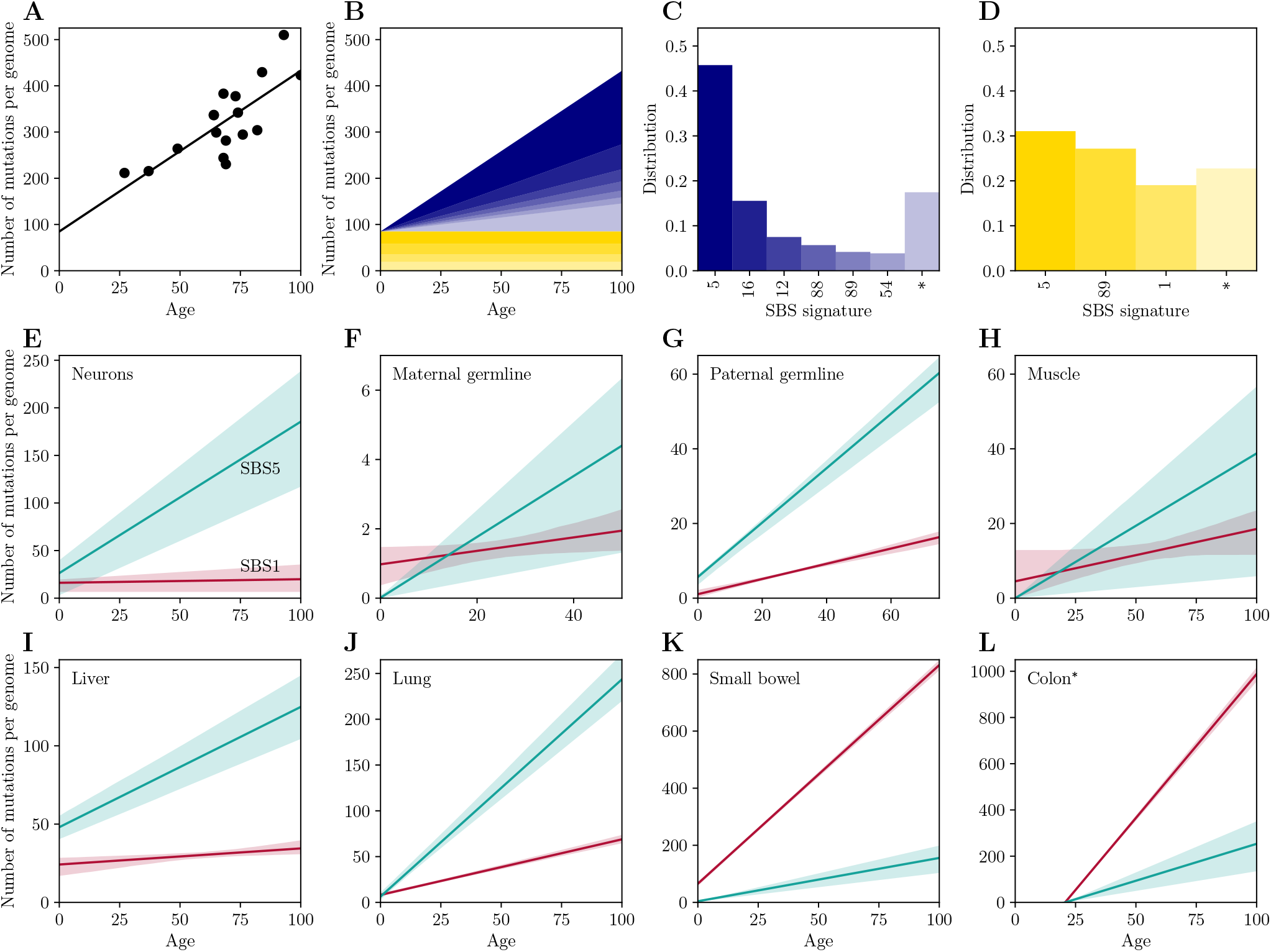
Clock-like mutational signatures in different cell types. (A) Age-dependent signature attribution of mutations in neurons. Shown is the increase in the number of mutations with age (reported per haploid genome); each point corresponds to a single donor. (B) The relative contributions of different signatures vary with age. The decomposition of the mutation spectrum into age-dependent (blue, C) and constant signature distributions (yellow, D). Mutational signatures are indicated by their COSMIC label; asterisks indicate unattributed signatures (see Methods). (E-L) The number of mutations assigned to clock-like signatures, SBS1 (red) and SBS5 (turquoise) in: (E) neurons, (F) maternal germline mutations, (G) paternal germline mutations, (H) smooth muscle from bladder, (I) liver hepatocytes, (J) lung epithelium, (K) small bowel epithelium, (L) colon epithelium (^∗^for this dataset, in which the intercept is negative, we assume both signatures increase with age). Throughout (E-L), shaded areas represent 95% confidence intervals, estimated by bootstrapping (see Methods).

In all cell types except for colonic epithelium, we find significant positive intercepts in the regression of the number of mutations on age, consistent with a burst of mutations in early development, for example. In these cases, we decompose mutations into the age-dependent distribution *P*_*a*_ and the constant component distribution *P*_*b*_. In the case of colonic epithelium (Figure 2L), the intercept is significantly negative, possibly because the mutation rate during ontogenesis is lower than in adult life (Figure A2B), when the exposure to damage and the cell division rate is higher [38]. To proceed with our analysis in this case, we assume (rather than infer, as for other cell types) that all signatures increase at constant rates with age.

Our analysis reveals multiple signatures that increase with age (see Figure A3), including the two that were previously reported [39], SBS1 and SBS5, and two additional signatures that are common across cell types considered, SBS12 and SBS16. SBS16 may not be independent from SBS5 [8, 10]. In turn, SBS12 contributes up to 7 −10% of mutations in neurons and the female germline, ~ 7% of mutations in lung and liver, ∼3% in small bowel and colon, ∼1% in the paternal germline, but is not found at detectable levels in muscle. Both SBS12 and SBS16 are of unknown etiology and dominated by T to C / A to G transitions.

In examining the mutation accumulation across cell types that vary in their division rates, we focus on SBS1 and SBS5, the two ubiquitous clock-like signatures, and use our decomposition method to examine possible sources for their age dependencies. If driven by cell division, we predict that the rate at which they will accumulate should vary substantially with cellular turnover rates. In contrast, if driven by damage rates, the rate should be much less sensitive to turnover rates, but may vary among tissues owing to differences in endogenous and exogenous damage rates.

In this regard, the accumulation of mutations with age observed in neurons is particularly informative, given that neurons are fully post-mitotic cells. Despite the lack of cell divisions, mutations accumulate at rates similar to actively dividing lineages [3]. Using our decomposition, signatures whose mutation numbers increase with age are distinct from those that do not (Figure 2C-D). Notably, the increase with age is predominantly driven by mutations assigned to signature SBS5, with secondary contributions from SBS16 and SBS12. Strikingly, there is no discernible contribution of SBS1, as we discuss in more detail below. In turn, mutations in the constant component are attributed primarily to signatures SBS5 and SBS1, as well as signature SBS89, which is of unknown etiology but has been reported to be active in the first decade of life [38].

Mutations assigned to the clock-like signature SBS5 are found across cell types and increase significantly with age in every one (Figure 2E-L). Moreover, SBS5 is the prevalent mutation signature in all cell types, except for small bowel and colon, for which more mutations are attributed to SBS1. That SBS5 is the dominant signature in post-mitotic cells such as neurons, as well as in maternal mutations, most of which arose in oocytes, indicates that such mutations are independent of DNA replication cycles and points to errors in DNA repair, which accumulate with damage rates (Eq. 1). Similarly, SBS12 and SBS16 contribute to both neurons and female germline mutations as well as to mutations in rapidly-dividing cells (Figure A3), suggesting that the age dependencies of these signatures is not driven by DNA replication cycles either.

When not driven by replication errors, our model predicts that the number of mutations will track the number of lesions and be clock-like so long as the product of the damage rate (*u*) and the incorrect repair rate (*ϵ*) is constant. If we assume that *ϵ* is fixed, as seems sensible if it is primarily determined by inherent properties of DNA repair proteins (e.g., the error rate of a polymerase), then the variation in the rate of SBS5 mutation across cell types reflects differences in rates of endogenous damage. Consistent with this notion, the rate of SBS5 mutation is highest for epithelia in the colon and lung (Figure 2), which plausibly experience high rates of damage, and lowest for mutations assigned to the maternal genome, potentially reflecting the fact that oocytes are particularly well protected [40, 41]. In some cases, SBS5 mutation accumulation may also result from an additional contribution of exogenous damage, helping to explain, for instance, the observation that increasing the damage rate by exogenous factors, such as long-term exposure to tobacco, significantly increases SBS5 mutation rate in lung cells [22, 42].

In that light, it may seem puzzling that in such different cell types, which presumably experience distinct sources of damage, a large fraction of mutations are consistently comprised of SBS5. As an explanation, we propose that SBS5 reflects errors in long-patch repair, a common repair pathway for many sources of damage [4]. Long-patch repair proceeds by excising multiple nucleotides surrounding the lesion and resynthesizing the gap using the intact strand. Since the errors of the gap-filling polymerase are often displaced from the position of the original lesion, the resulting mutations should reflect the error profile of the polymerase, regardless of the molecular signature of the original damage process.

The second signature to increase with age, SBS1, does so in all cell types considered, except for liver, where hepatocytes are routinely dormant in the cell cycle [43], and neurons, which are post-mitotic. More generally, the rate at which mutations assigned to SBS1 increase with age varies widely among cell types and is highest in those characterized by the highest turnover rates (such as intestinal epithelia, where turnover time estimates are of the order of 3 days [44]). Thus, SBS1 appears to be driven by cell division rates. A possible exception is the observation of a slight increase with age in maternal germline mutations, most of which arose in oocytes (see the discussion of germline mutations below). These observations are in agreement with previous observations from cancer studies [6, 39, 45] and indicate that, while it would seem intuitive that mutations driven by deamination—a spontaneous, ongoing process—should accumulate with absolute time, instead they track cell divisions.

Mutations that arise from damage will track cell division if efficiently and accurately repaired (Figure 1). Current data do not allow us to pinpoint when in the cell cycle the damage accrues, however. Two plausible sources are unrepaired mismatches that accrue during the cell cycle and deamination of single-stranded cytosines during DNA replication. We model their possible contributions in Appendix A; as we show, their relative importance will depend on efficiency of mismatch repair as well as the length of time spent singlestranded during replication, parameters that are to our knowledge unknown.

Regardless, we can use the fact that SBS1 does not discernibly increase with age in post-mitotic neurons in order to estimate an upper bound on the rate of SBS1 mutations due to repair errors. Given no cell divisions, *ϕ* = 0, we estimate the upper limit of the error rate of the resolution of T:G mismatches at CpG sites to be 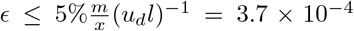 (here, we assume the detection threshold to be ≤5%, the lowest loading of an attributed signature in neurons). Transient single-strandedness due to transcription would enhance the rate of deamination [46], in which case the upper bound on the error rate of repair would be lower. Our estimate is similar to the measurements of the fidelity of polymerase beta [47], employed by base excision repair (BER), one of the pathways that repairs T:G mismatches [48]. It is also of the order of the lower bound on the error rate for DNA synthesis without proofreading, *ϵ*_0_ ∼10^−4^, found by considering the equilibrium kinetics of DNA synthesis, given the energy difference between a mismatch and a correct base pair [49, 50]. These calculations show that, despite a high rate of spontaneous deamination, repair mechanisms should be accurate enough for incorrect repair to be an insignificant source of mutations. Instead, repair of this type of mutation is likely both very efficient and accurate in all cell types, leading the number of SBS1 mutations to track cell divisions.

### Mutation accumulation in the germline

While SBS1 and SBS5 are known to predominate among germline mutations [12, 33], we expect the rates of mutation accumulation to differ between the sexes, given the pronounced differences in gametogenesis. In mothers, any mutation that increases with maternal age should have arisen in an oocyte, a post-mitotic cell (although a small fraction may also arise in the early development of the child, if children of older mothers have more mutations in the first few cell divisions [51]), whereas in fathers, age-dependent mutations should arise in dividing spermatogonia. In turn, the mutations that do not depend on parental ages in either sex originated either prior to the onset of puberty in the parents or soon after fertilization of the offspring [52]. Given that in humans, early development is the same in both sexes until the ∼6th week of embryonic life [53], we might expect some similarity between the mutation types that contribute to the constant component in the two sexes.

In agreement with these expectations, for maternal mutations, the distributions of mutational signatures differ markedly between the age-dependent and constant components: in the constant component distribution (Figure 3D and 3H), C to T / G to A transitions dominate, with leading contributions of SBS1, SBS6 (associated with defective DNA mismatch repair [8]), and SBS30 (associated with defective base excision repair [54]). The age-dependent distribution is significantly more diffuse (Figure 3C and 3G), with top contributions from SBS5, SBS12, SBS16, and SBS39. SBS39 predominantly features C to G / G to C transversions, a substitution type known to increase sharply with maternal age and associated with double strand break repair [51, 55]. Given that mutational signatures have been identified from cancer somatic tissues [8], it is conceivable that SBS39 absorbs a significant portion of the C to G / G to C substitutions characteristic of maternal mutations, even if the process that generated them in the germline has a distinct etiology. Qualitative conclusions are similar when adding the father’s age as a covariate in the model (see Figure A4 in the Appendix).

**Figure 3.**
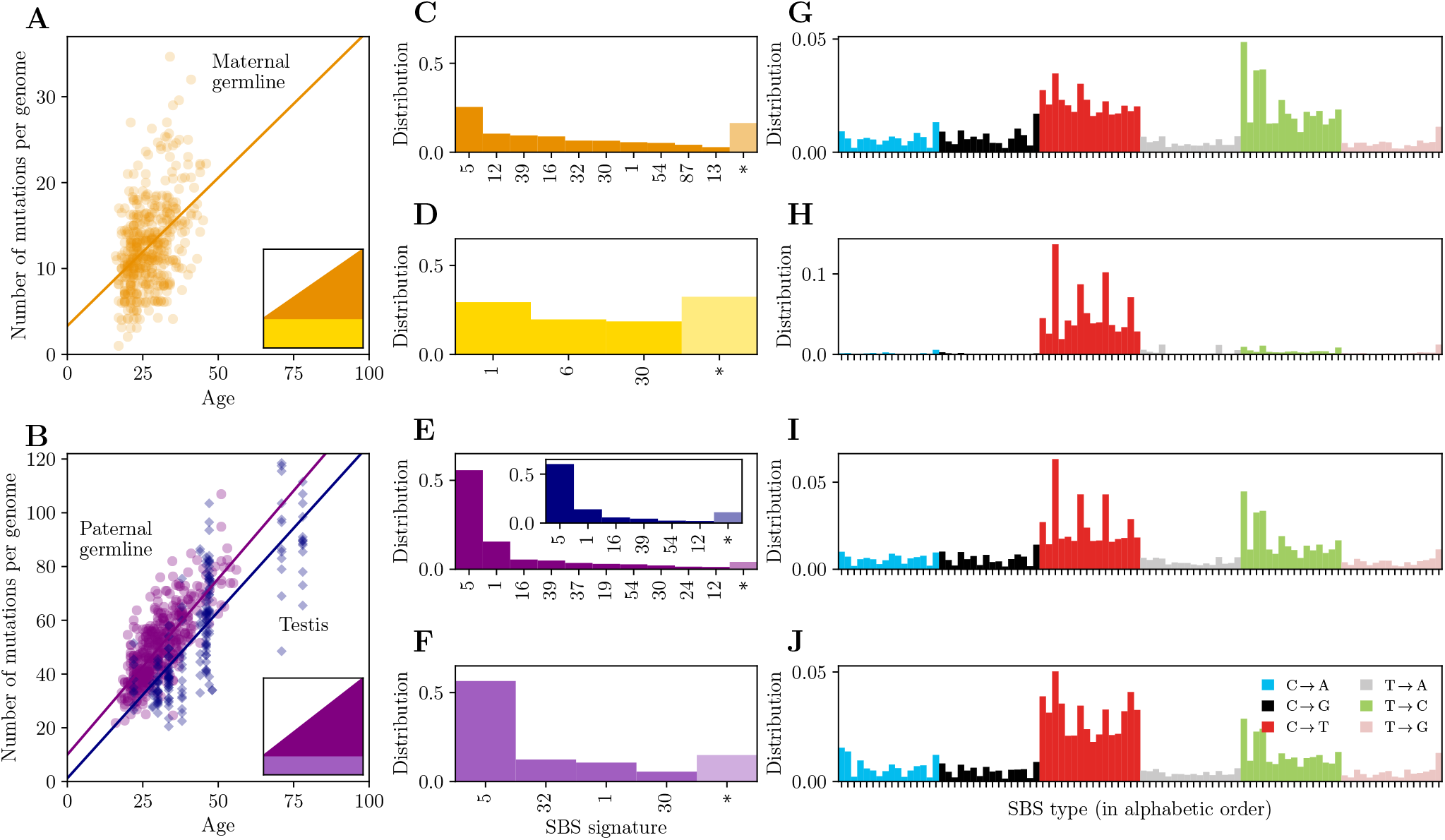
Signature attribution for germline mutations. (A) Effect of maternal age on mutations assigned to the maternal germline in pedigree data. (B) Effect of paternal age on mutations assigned to the paternal germline in pedigree data. Paternal germline mutations (purple) accumulate at similar to rate to somatic mutations in testis seminiferous tubules (blue). (C-F) The decomposition of the mutation spectrum into age-dependent (C,E) and constant signatures (D,F) in maternal and paternal mutations; see (A) and (B) for the color code. The inset in (E) shows the decomposition of somatic mutations in the testis seminiferous tubules. SBS signatures are indicated by their COSMIC label; asterisks indicate the contribution of unattributed mutations (see Methods). (G-J) The distribution of SBS types reconstructed using the signature attributions (C-F). SBS types are grouped by the six substitution types (see J for color code) and sorted alphabetically by the sequence context (ACA to TTT).

Surprisingly, there is a small but significant increase of SBS1 mutations with maternal age (0.019 mutations per year per gamete), which seems at odds with the lack of cell divisions in oocytes. One possibility is that there is an increase in the steady state number of T:G mismatches in aging oocytes, potentially due to reductions in repair efficiency *r* in older mothers [51]. Uncorrected T:G mismatches in oocytes would lead to zygotic mutations in the first division after fertilization and be detected by pedigree sequencing of trios. Alternatively, the slight increase of SBS1 mutations with age in maternal mutations, in contrast to what is seen in neurons, could be explained by a higher error rate of mismatch repair in oocytes. In other words, while the rate of erroneous repair of T:G mismatches due to cytosine denomination may be minimal in neurons, it could be higher and detectable in oocytes. In principle, a similar outcome would be observed if the rate of erroneous repair is the same but cytosines deaminate more often in oocytes relative to neurons, but this explanation seems less likely given the low DNA methylation levels in maternal gametes [56].

Among paternal mutations, the age-dependent and age-independent distributions are both dominated by signature SBS5. Most of the mutations in the constant component (Figure 3F) are attributed to SBS5, but there is also an enrichment of C to T / G to A transitions (SBS32, SBS1, and SBS30). Unlike in mothers, the increase with age is driven by SBS5 and SBS1 jointly (Figure 3E). To test whether age-dependent paternal germline mutations accumulate primarily in spermatogonia, as expected, we compare these findings with the decomposition of somatic mutations detected in the seminiferous tubules of the testes [2]. As reported by [2], the slope of the regression line is very similar for the two datasets (Figure 3B). In contrast to their study, however, the intercept for testis is not significantly different from 0, while the intercept for germline mutations is of ~ 10 mutations per haploid genome; the reasons for this difference are unclear to us. Regardless, in our analysis, the age-dependent component of paternal mutations has a very similar distribution of signatures to those inferred for testis (Figure 3E and inset).

Overall, the age-dependent distribution is highly similar in the two sexes (Figure 3G and 3I), except for SBS1, which is significantly more pronounced in the paternal germline. The similarity of age-dependent mutational signatures between the two sexes suggests that it is not only the continuous cell divisions but also the higher damage rate that distinguishes the paternal germline from the maternal one. While the constant components differ between maternal and paternal mutations, SBS1 and SBS30 are found in both, likely reflecting a contribution of gonosomal mutations.

## III. DISCUSSION

As we show by modeling, distinct mutagenic processes can give rise to the clock-like accumulation of mutations observed in the germline and soma, so long as cell divisions and damage occur at a reasonably constant rates. To tease these processes apart, we estimate the rate of accrual of clock-like mutations across dividing and non-dividing cell types. Our analysis reveals that the ubiquitous SBS1 and SBS5 originate predominantly from different sources: whereas SBS1 tracks cell divisions, SBS5 accumulates in post-mitotic as well as dividing cells, and appears to track DNA damage levels.

Based on the behavior of SBS5 across cell types, we hypothesize that such mutations arise from errors in long-patch DNA repair. This hypothesis could be explored further by examining, for example, if SBS5 mutations are enriched at loci with high repair rates, or if they tend to co-occur within distances consistent with the action of long-patch repair. In turn, the dependence of SBS1 on cell divisions suggests that the rate of accumulation of such mutations could serve as a “counter” for cell divisions [39, 57], applicable to different cell lineages. For now, however, cell division rates at different stages of human development remain poorly characterized, stymieing such efforts.

Our analyses further confirm the importance of damage as a source not only of somatic mutations, but also for germline mutations [51, 58, 59]: over two-thirds of germline mutations are assigned to SBS1 and SBS5, signatures that arise from damage that is either not repaired or repaired incorrectly. The source of such DNA damage is most likely endogenous cellular processes, accounting for the omnipresence of these two signatures across cell types [2] and species [60], as well as their characteristic clock-like behavior under most conditions [42].

While our analyses help to make sense of a number of disparate observations in human germline and soma, they also raise a number of new questions. In particular, it is puzzling that CpG transitions accumulate with absolute time in phylogenetic data from mammals [61, 62], when SBS1 mutations accumulate with cell divisions within humans, and there are dramatic differences in germ cell division rates across mammalian species. It is also unclear how species-specific rates are set for other types of mutations. The dominance of SBS5 in the germline implies that most of the variation in mutation rates across species is likely explained by differences in the rates of DNA damage and repair (e.g., [63, 64]). To what extent this balance is directly shaped by selection versus a byproduct of changes in cellular activity remains to be explored, however.

## Iv. METHODS

### Model of mutation accumulation

We first consider the interplay of damage and repair during the cell cycle. The time since the last DNA replication is denoted by *t*. We denote the number of intact sites by *n*_0_, the number of lesions by *n*_1_, the number of mismatches by *n*_2_, and the number of mutations by *n*_3_, where all these sum up to the genome size, i.e., *n*_0_ + *n*_1_ + *n*_2_ + *n*_3_ = *l*. DNA damage leads to new lesions at rate *u*. Lesions are repaired at rate *r*, with error rate *ϵ*. Incorrectly repaired lesions lead to mismatches, which are resolved at rate *q*. We assume the mismatch resolution mechanism cannot discern the correct base in the mismatch site. We further assume for simplicity that this process is independent of the molecular context of the mismatch and thus leads to a mutation with probability 0.5 (see schema in Figure 4). The model parameters and variables are summarized in Table I.

**Figure 4.**
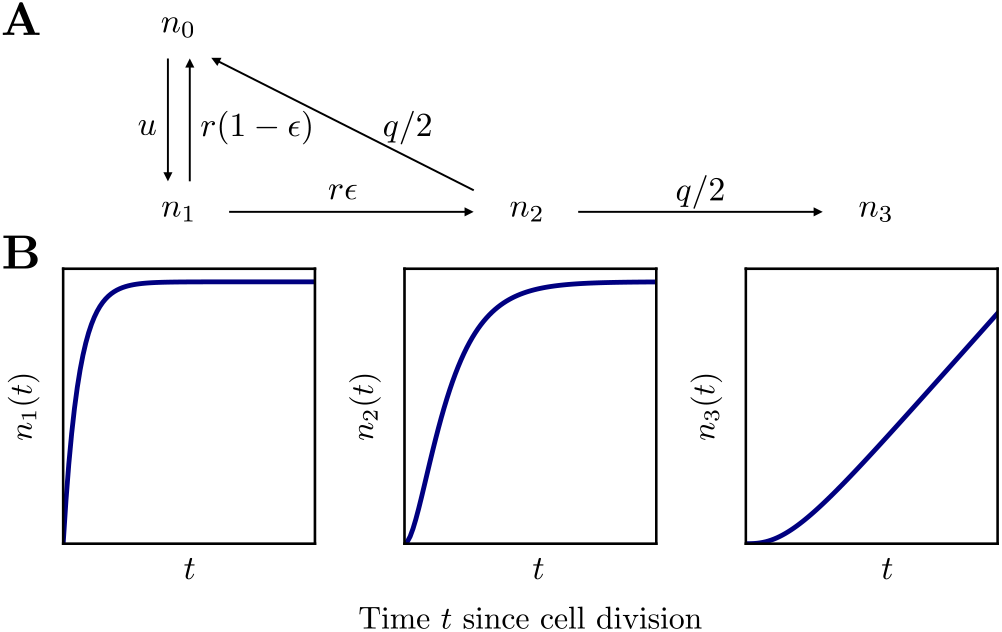
(A) Kinetics of the interplay of damage and repair. The two-state model used for CpG transitions accumulation outlined in Appendix A is obtained by taking the limit *q* → 0 (see below). (B) Example solution of the model for the number of lesions (left), mismatches (center), and mutations (right).

**Table I.** Parameters and variables of the model.

| Notation | Parameter / variable |
| --- | --- |
| $l$ | Genome size |
| $n_0$ | Number of intact sites |
| $n_1$ | Number of lesions |
| $n_2$ | Number of mismatches |
| $n_3$ | Number of mutations |
| $u$ | Damage rate |
| $r$ | Repair rate |
| $\epsilon$ | Repair error rate |
| $q$ | Mismatch resolution rate |
| $\phi$ | Cell division rate |

The expected number of sites in each state is well approximated by the following system of equations

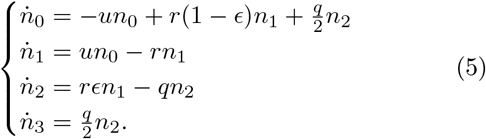

where we have neglected the probability of damage affecting the same site multiple times, as is realistic. The number of intact sites is always much greater than the number of lesions, mismatches and mutations combined, i.e., *n*_0_ ≈*l* ≫*n*_1_ + *n*_2_ + *n*_3_. This dynamics is therefore well approximated by

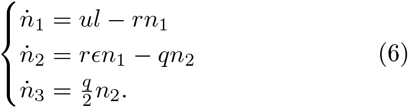

The solution to this system with initial condition *n*_0_(0) = *l* (and thus *n*_1_(0) = *n*_2_(0) = *n*_3_(0) = 0) is

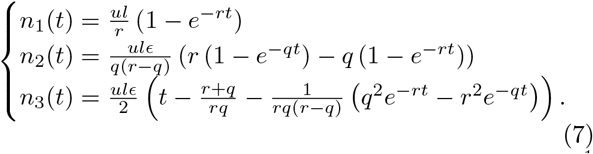

At steady state, which is achieved at time *t* ≫*r*^−1^ and *q*^−1^, the expected numbers of lesions and mismatches are constant, and the number of mutations *n*_3_ grows linearly with time. Specifically,

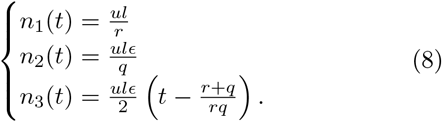

where 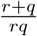 is the delay due to lesion and mismatch processing. We assume, as is plausible, that a steady state is established before cell division, which in terms of model parameters means that the rate of cell division obeys *ϕ* ≪ *r* and *q*, and consequently 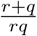 is negligible at timescales of *ϕ*^−1^ or longer.

Next, we consider the expected number of mutations *m* over a period *x* that spans multiple cell divisions occurring at rate *ϕ*. We additionally account for unrepaired replicatory errors, of which there are *wlP* per cell division, where *w* denotes the error rate of a replicative polymerase and *P* is the probability that a mismatch is unrepaired by mismatch repair. Altogether, the expected number of mutations is given by

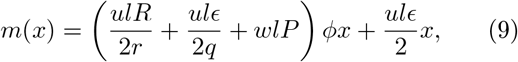

where we assume that unrepaired lesion leads to a mutation in one of the two dividing cells with probability *R*.

To quantify variation in the number of mutations, we note that the set of equations (6) corresponds to a monomolecular reaction system and therefore that the equivalent chemical master equation is solved by a product Poisson distribution [65]. Thus the variances of the random variables *N*_1_, *N*_2_, and *N*_3_ (numbers of lesions, mismatches, and mutations) are equal to their expected values and given by

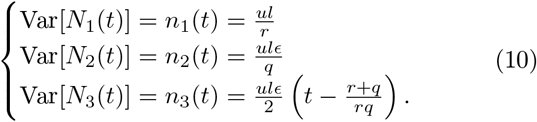

We further assume that cell division dynamics is independent of the dynamics within a cell cycle, and that the number of cell divisions in time *x* is Poisson distributed with mean *ϕx*. We can therefore estimate the variance of the number of mutations *M* at age *x* using the law of total variance,

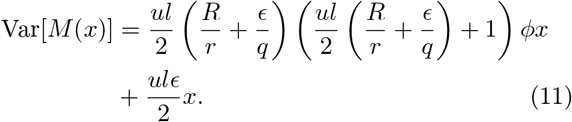

To obtain the reduced model describing the dynamics underlying CpG transitions (see Figure A1 in Appendix A), we take the limit *q* → 0. This results in

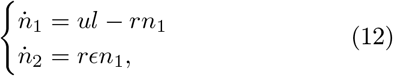

where *n*_1_ now stands for the number of T:G mismatches, and *n*_2_ is the number of mutations. This system is solved by

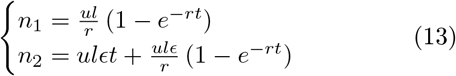

Unrepaired mismatches lead to a mutation in one of the two dividing cells. Additionally, we consider that cytosines can deaminate at rate *u*_*s*_ during the transient single-strandedness that occurs at DNA replication, for time ∆*t* (see Appendix A for details). Analogously to the general model, assuming the dynamics reaches a steady state before the cell division, the expected number of mutation is given by

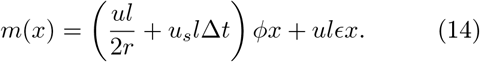

### Data analysis

We analyze the patterns of mutation accumulation in different cell types, using a variety of publicly available data sets that each include seven or more individuals of different ages. The studies used a variety of approaches in order to characterize mutations that accrue in a given cell type, including single molecule sequencing (S), sequencing of small monoclonal samples (C),and sequencing of cell colonies derived from a single cell (D). In addition, we consider *de novo* mutation calls based on sequenced genomes from blood samples of human trios (P). In this case, mutations are assigned to maternal and paternal genomes using “read-backed phasing” [55, 66] as well as by transmission to a third generation[37, 55]; such mutations are assumed to have arisen in the oocyte and during spermatogenesis, although there is also a small contribution of gonosomal mutations and mutations that arose in the early development of the child [37]. We only select probands in which the parental origin has been determined for over 90% of germline mutation calls. The data set (see Table II below) includes cell types with varying cell division rates, ranging from post-mitotic neurons to rapidly dividing epithelial cells from small bowel and colon [67]. We excluded data from donors with cancers in the focal tissue, and past and current smokers from the lung data set [22]. Download links to the data are provided in a Supplementary Table.

**Table II.**
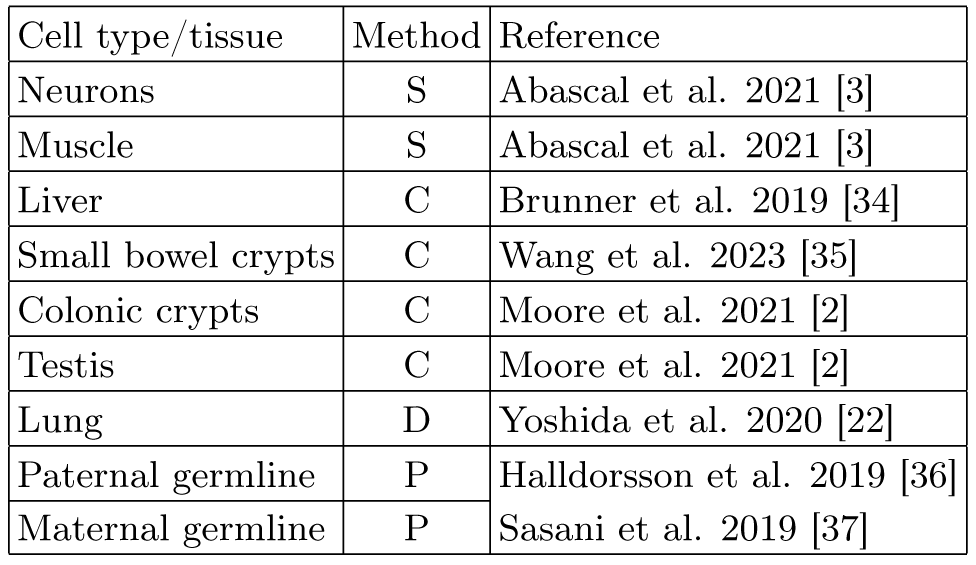
Datasets used in the analysis of clock-like mutation accumulation.

We annotated each SBS substitution with its local context, i.e., the 5’ and 3’ flanking nucleotides, using the GRCh37 genome reference [68]. Using this annotation, we classified each substitution into one of the 192 types (there are 4^3^ = 64 different 3-mer contexts, and for each of them, there are three possible substitutions). This classification was then collapsed to give the standard 96 categories, which are strand-invariant [6, 7].

### Mutational signatures attribution

First, we describe how we decompose a set of mutations as a linear combination of 79 COSMIC version 3.3 SBS mutational signatures [10]. For any mutation of SBS type *z*, the probability that it is attributed to a given signature class *s* is given by Bayes’ formula

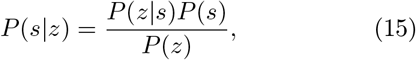

where *P* (*z*) = Σ_*s*_ *P* (*z*| *s*)*P* (*s*). *P* (*z*| *s*) is known, but we need to infer the set of loadings P = *P* (*s*) from the data.

The log-likelihood of the loadings P given the observed mutations 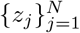 is given by

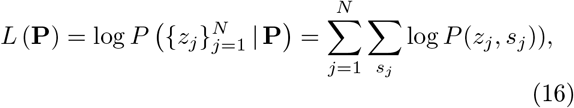

where *P* (*z*_*j*_, *s*_*j*_) is the joint probability of mutation of type *z*_*j*_ and the underlying signature *s*_*j*_ (a hidden variable). We maximize the likelihood iteratively using the expectation-maximization method (EM, [69]), under the constraint of normalization, _*s*_ *P* (*s*) = 1, and *P* (*s*) ≥0 for all *s*.

### Initial condition

We initialize the optimization with a uniform distribution.

#### E step

Provided the distribution at iteration *i*, P^*i*^ = {*P* ^*i*^(*s*)}, we compute the pseudo-log-likelihood of the set of loadings P,

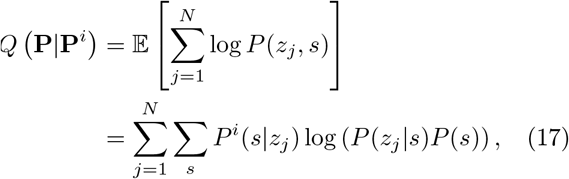

where

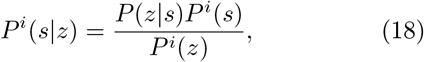

and *P* ^*i*^(*z*) = Σ_*s*_ *P* (*z*|*s*)*P* ^*i*^(*s*).

#### M step

We find the distribution in the next iteration, P^*i*+1^, by maximizing the pseudo-log-likelihood *Q*(P|P^*i*^) under the constraint *P* (*s*) = 1, i.e.,

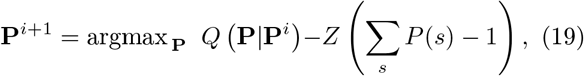

where *Z* is a Lagrange multiplier. In this way, we find that

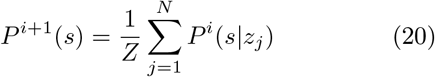

and the value of the multiplier can be found by applying the condition of normalization, 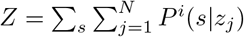

We apply regularization to avoid overfitting. Specifically, we impose sparsity on the distribution P^*i*^ by introducing a Dirichlet prior over the set of distributions P,

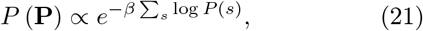

where *β* ≥0 parametrizes the degree of sparsity. We discuss the choice of regularization strength below.

With this regularization, the distribution P^*i*+1^ in the next iteration takes a closed form. Taking advantage of the fact that a Dirichlet prior is conjugate to multinomial likelihood (16), it can be shown [70] that

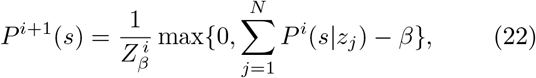

where the normalization factor reads 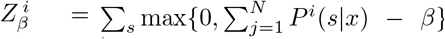 . This expression implies that if the estimated loading of a signature *s* is too low, it is set to 0.

### Convergence criterion

We iterate until convergence, i.e., when for all signatures *s* we have | |*P* ^*i*+1^(*s*) − *P* ^*i*^(*s*) | | *<* 10^−4^.

In what follows, we extend this signature attribution method to infer age-dependent signatures in data consisting in mutations from donors of varying ages. First, we preform a linear regression on the number of mutations per haploid genome, *y*, on age, *x*

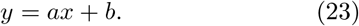

We find *a* and *b* for each cell type, using the scipy package [71], see Figure A5. Next, we assign the mutational signatures that make up the slope and the intercept. Namely, we assume that independent distributions govern the signature compositions of the age-dependent and age-independent mutations and denote them P_**a**_ = *P*_*a*_(*s*) and P_**b**_ = *P*_*b*_(*s*), respectively. The expected proportion of mutations of type *z* at age *x* is then given by

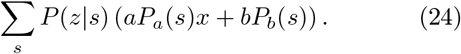

where we sum over COSMIC SBS signatures.

In order to pose the problem of estimating the two components, P_**a**_ and P_**b**_, we assign to two hidden variables to each mutation *j*: the signature to which the mutation is attributed, *s*_*j*_, and an indicator variable *I*_*j*_, which equals 1 for age-dependent mutations and 0 for age-independent mutations. In these terms, the loglikelihood of the data given the two sets of loadings is

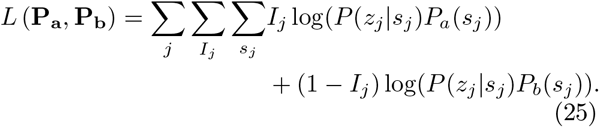

We estimate the loadings using expectationmaximization, as previously described. In this case, the pseudo-log-likelihood is

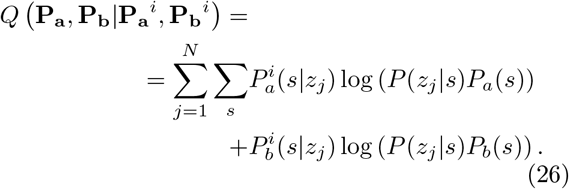

where the two sets of membership probabilities 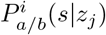 are computed analogously to Eq. (18).

We assess the uncertainty of our estimates by bootstrapping. For each data set, we resample cells (or trios in the case of pedigree data) with replacement 1000 times. For each replicate, we estimate the slope and intercept of the age dependence and infer the corresponding signature attribution. We test for the significant presence of a given signature by performing the inference with weak regularization (i.e., with *β* = 0.1). This choice of regularization strength does not affect the main components and only zeroes out the components of the decomposition with very low membership probabilities (see Figure A6), as specified by Eq. 22. We include a signature *s* from a given data set if the probability *P*_*a/b*_(*s*) is nonzero in at least 95% of bootstrap replicates; in the figures, we denote the residual contribution of signatures that do not meet this condition with an asterisk.

## Acknowledgements

We thank Ziyue Gao and members of the Andolfatto, Przeworski, and Sella labs for useful discussions. This work was supported by HFSP postdoctoral fellowship LT000257 to MdM and R01 GM83098 to MP.

## APPENDIX

### A. Model of CpG transitions accumulation

The process of spontaneous cytosine deamination contributes substantially to the mutation rate [17]. The standard explanation is that because at methylated cytosines, the deamination rate results in a thymine, one of the canonical bases, the efficiency of repair is low [72]. To investigate the dynamics underlying the accumulation of CpG transitions, we therefore consider the consequences of methylated cytosine deamination during the cell cycle. Given that deamination leads directly to a mismatch, we can employ a simpler model, following [24]. The dynamics is analogous to the general model described in the main text and can be recovered in the *q* → 0 limit (see Methods, Eq. 12).

Methylated cytosines in double-stranded DNA deaminate spontaneously at a high rate (estimated as *u*_*d*_ = 2.3 × 10^−5^ per year [46]). The resulting T:G mismatches can be detected and repaired by base excision repair (BER, [48]) or mismatch excision repair (MMR, [73]) at effective rate *r*. With probability *ϵ*, the repair machinery erroneously substitutes the guanine for an adenine, leading to a mutation (Figure A1A).

As in the more general model, at early times, *t < r*^−1^, the number of mismatches grows linearly with the rate *u*_*d*_, eventually reaching a steady state with the expected number of mismatches equal to *u*_*d*_*l/r*, where *l* stands for the total number of methylated cytosines. Given the high estimated rate of spontaneous deamination and the relatively low observed rates of SBS1 mutations [13], repair must be efficient and have time to act before the cell division. The implication is that the repair rate is higher than the cell division rate, *r* ≫*ϕ*, and that the cell divides after the number of mismatches has reached a steady state. As unrepaired mismatches lead to mutations in one of the two daughter cells, DNA replication results in mutation. Therefore, the number of mutations *m* at age *x* depends on the cell division rate *ϕ*. On the other hand, the number of mutations due to defective repair follows absolute time, and such mutations accumulate independently of cell divisions.

Under an alternative hypothesis that is not mutually exclusive (Figure A1B), CpG transitions result from the deamination of methylated cytosines in single-stranded DNA immediately prior to replication and cannot be repaired. The number of mutations that arise from such deamination events is proportional to the expected number of cell divisions *ϕx* and the time of transient single-strandedness ∆*t*. The rate of deamination of methylated cytosines in single-stranded DNA is estimated to be orders of magnitude higher than in doublestranded DNA (*u*_*s*_ = 3.5 ×10^−3^ per year [46]), making this scenario a plausible explanation for the observed

SBS1 mutation rates: to account for 1 mutation in 10 divisions, the time of transient single-strandedness would need to be of order ∆*t* ∼1 minute. This order of magnitude can be compared with the typical velocity of replication fork in human cells, of order 1 kb per minute [74].

It has also been proposed that some of the CpG transitions could reflect DNA replication errors, if methylated cytosines are a difficult template for DNA polymerases [75, 76]. Unlike other replication errors, however, T:G mismatches can be repaired by BER or MMR throughout the cell cycle and therefore, the contribution of this type of replication error to the SBS1 mutation rate may be negligible.

### B. Supplementary figures

**Figure A1.**
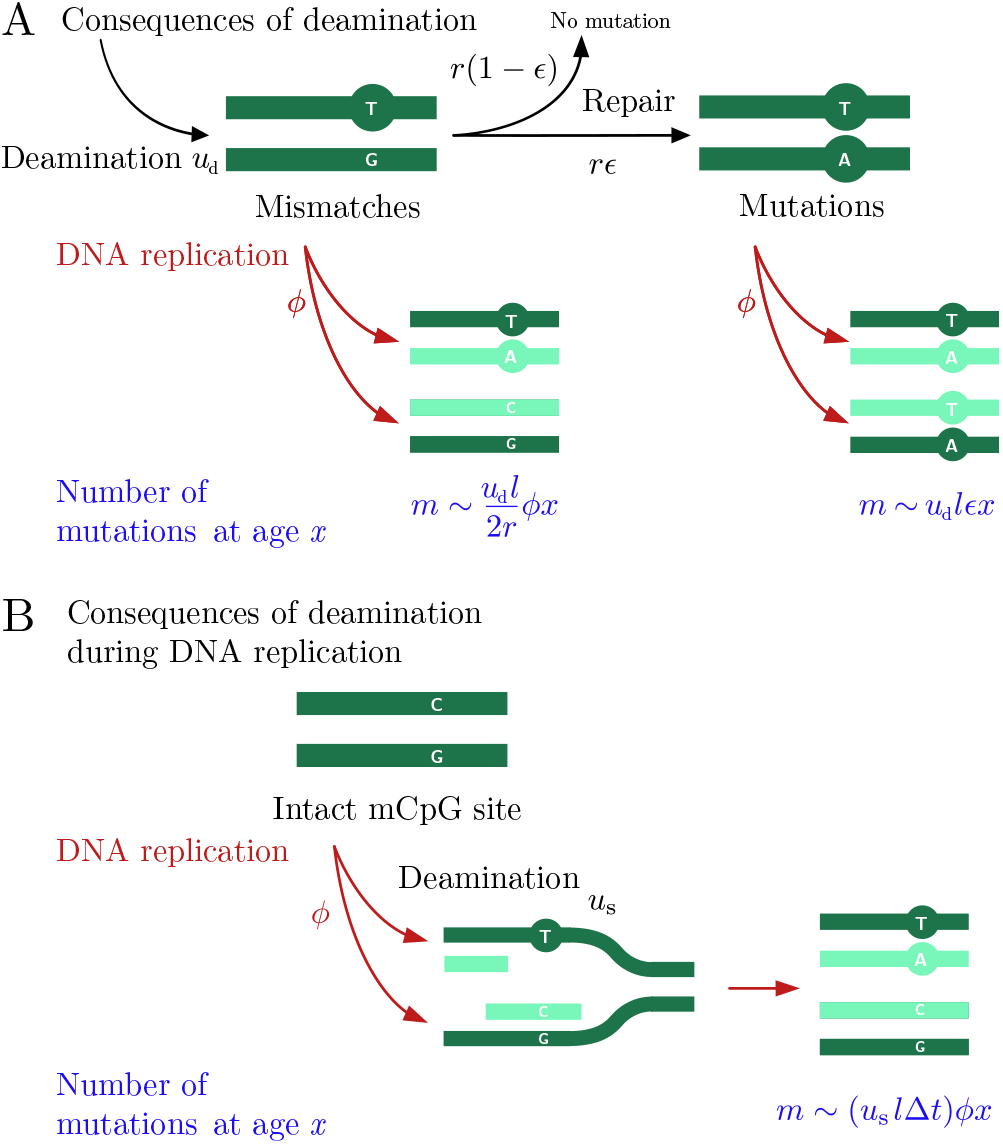
Model of the accumulation of CpG transitions. (A) Model of the consequences of spontaneous deamination of methylated cytosines during the entire cell cycle [24]. Double-stranded cytosines deaminate at a constant rate *ud* per basepair and the resulting mismatches are repaired at a rate *r*. With probability *ϵ*, the mismatch resolution is incorrect and leads to a mutation. We assume that the cell divides immediately after DNA replication and treat the two processes as simultaneous (occurring at a rate *ϕ*). Unresolved mismatches at cell division lead to a mutation in one of the two daughter cells. We provide the prediction of the model for the number of mutations *m* at a given age *x*. Inefficiently repaired mismatches accumulate with the cell divisions, which occur at rate *ϕ*, and repair errors accumulate with absolute time, independent of cell divisions. The number of methylated cytosines is denoted by *l*. (B) An alternative source of CpG transitions is the deamination of methylated cytosines in single-stranded DNA (at a rate of *us*) during DNA replication. The model assumes that a deamination immediately preceding the polymerization of the second strand is not repaired and leads to a mutation in one of the daughter cells. The expected number of mutations is proportional to the rate of cell divisions and the time of transient single-strandedness ∆*t*.

**Figure A2.**
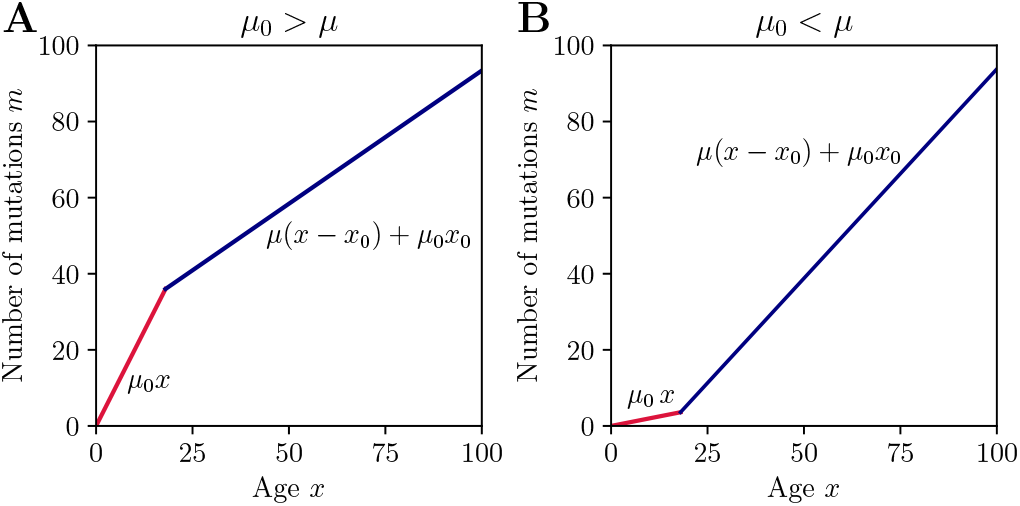
Piecewise linear model. Until age *x*_0_ = 18 mutations accumulate at rate *μ*_0_ (in red) and after age *x*_0_ at rate *μ* (blue). (A) When *μ*_0_ *> μ*, for data from donors of ages *x > x*_0_, a regression would yield a positive intercept, *b >* 0. (B) When *μ*_0_ *> μ*, such regression would yield a negative intercept, *b <* 0.

**Figure A3.**
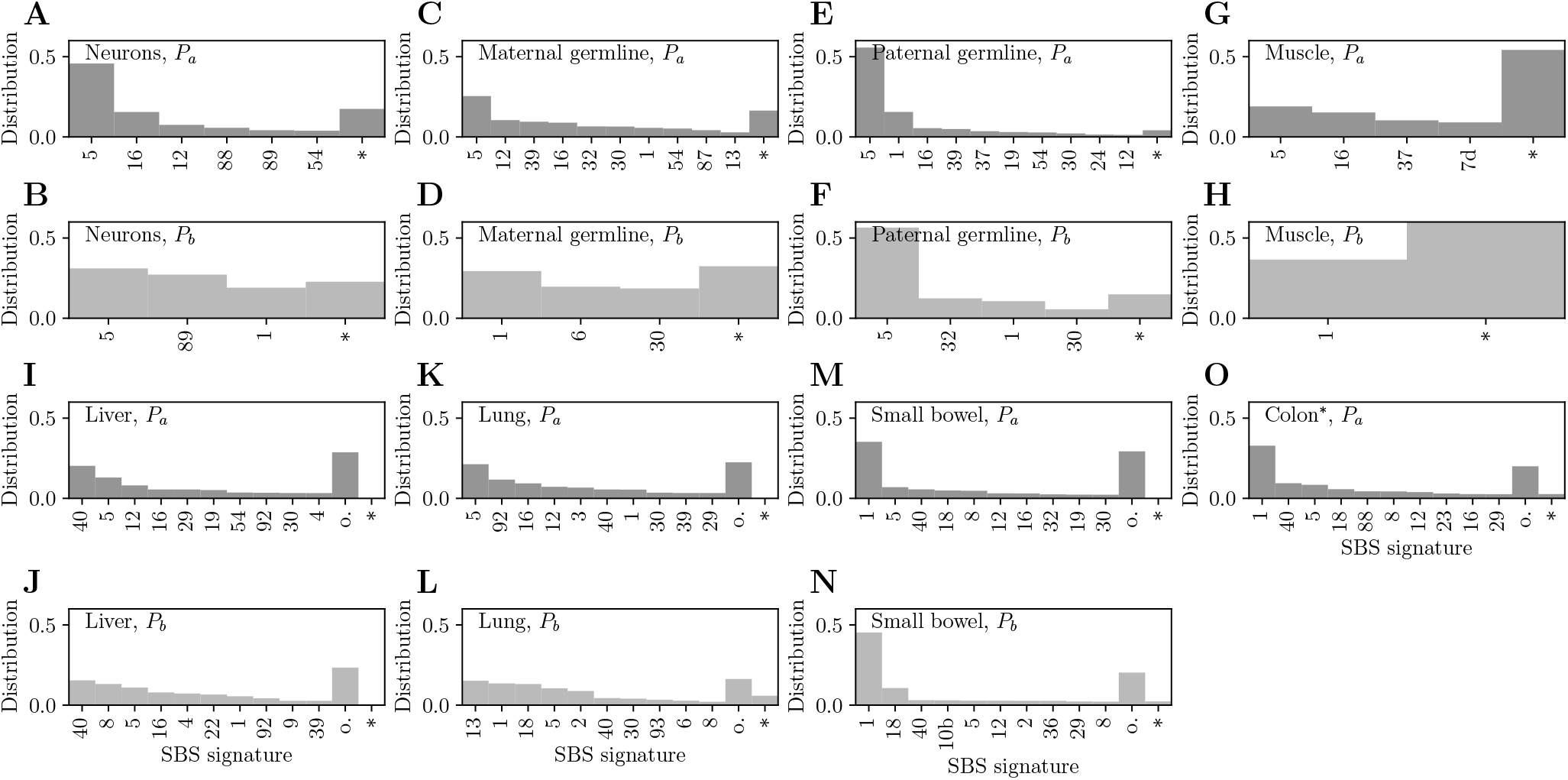
Inference results with weak regularization (*β* = 0.1): loadings for the age-dependent component, *P*_*a*_, and the constant component, *P*_*b*_, in (A,B) neurons, (C,D) maternal germline mutations, (E,F) paternal germline mutations, (G,H) smooth muscle from bladder, (I,J) liver hepatocytes, (K,L) lung epithelium, (M,N) small bowel epithelium, (O) colon epithelium (^∗^for this dataset, in which the intercept is negative, we assume all signatures increase with age). Signatures are indicated by their COSMIC label; we only show top 10 attributed signatures, and “o.” stands for other attributed signatures. An asterisk indicates the contribution of unattributed signatures (see Methods).

**Figure A4.**
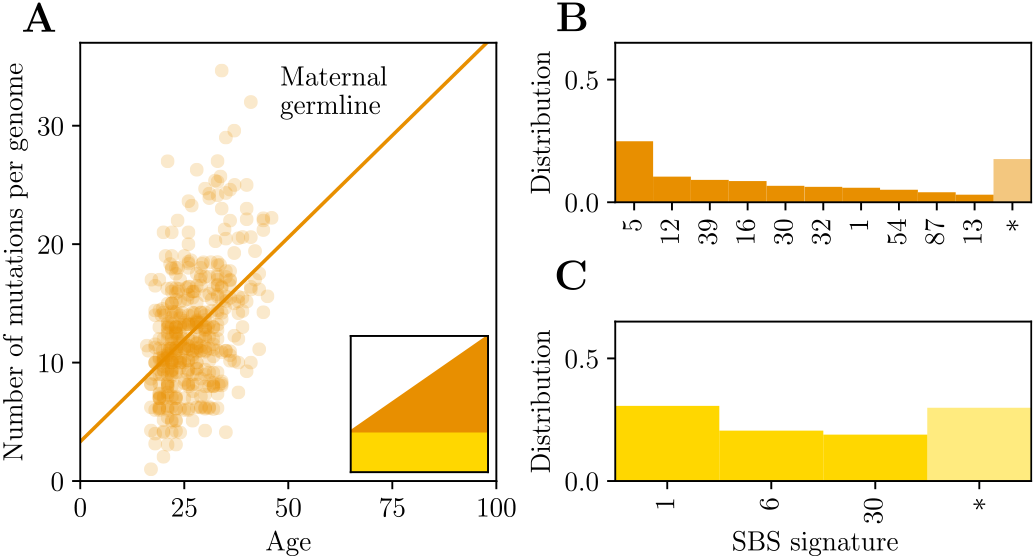
Age-dependent signature attribution for germline maternal mutations. (A) Effect of maternal age on mutations assigned to the maternal germline in pedigree data. Paternal age was included as a covariate in linear regression. (B-C) The decomposition of the maternal mutation spectrum into age-dependent (B) and constant signatures (C); see (A) for the color code. SBS signatures are indicated by their COSMIC label; asterisks indicate unattributed signatures (see Methods).

**Figure A5.**
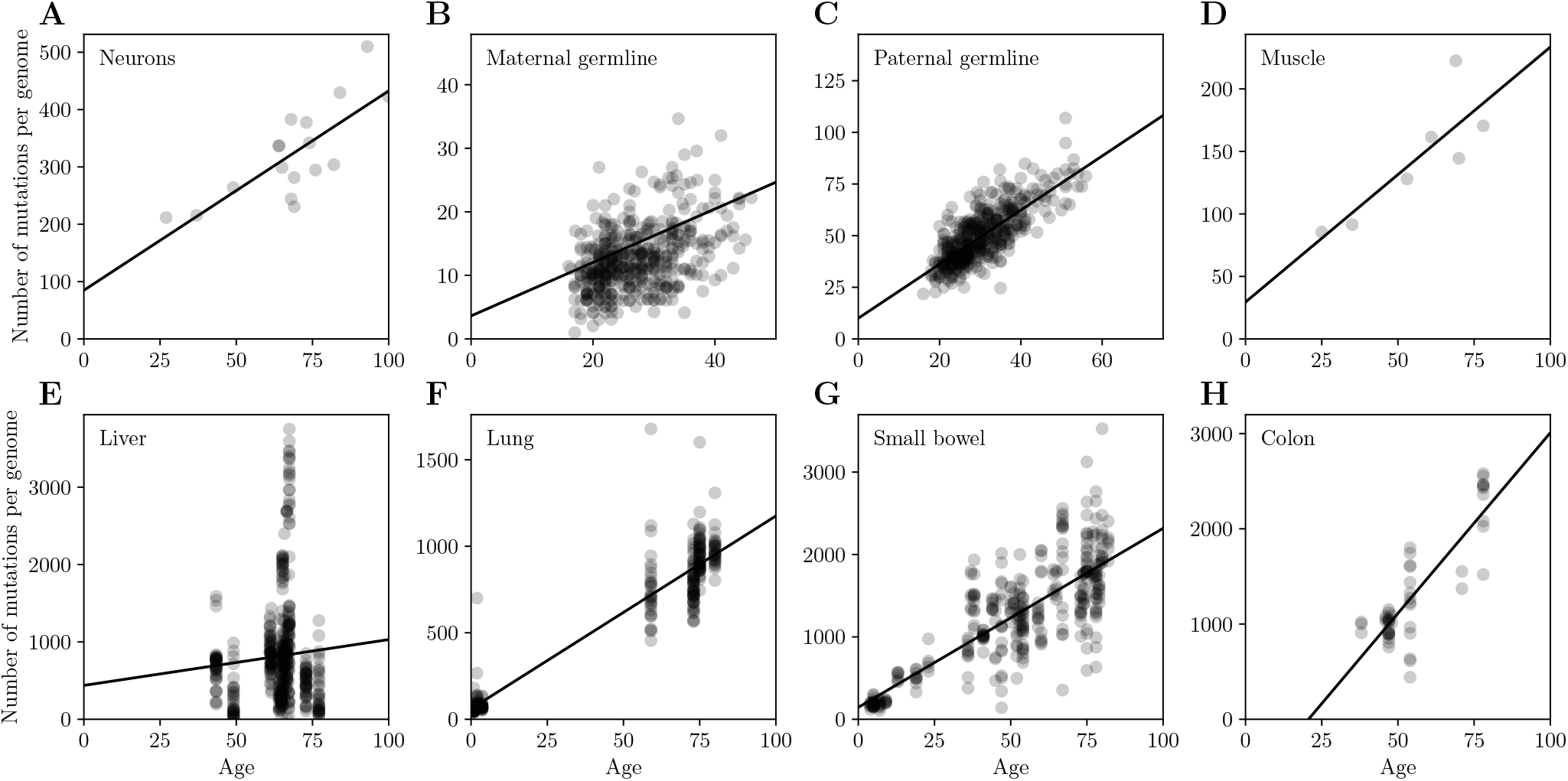
The total number of mutations as a function of age in: (A) neurons, (B) maternal germline mutations, (C) paternal germline mutations, (D) smooth muscle from bladder, (E) liver hepatocytes, (F) lung epithelium, (G) small bowel epithelium, (H) colon epithelium.

**Figure A6.**
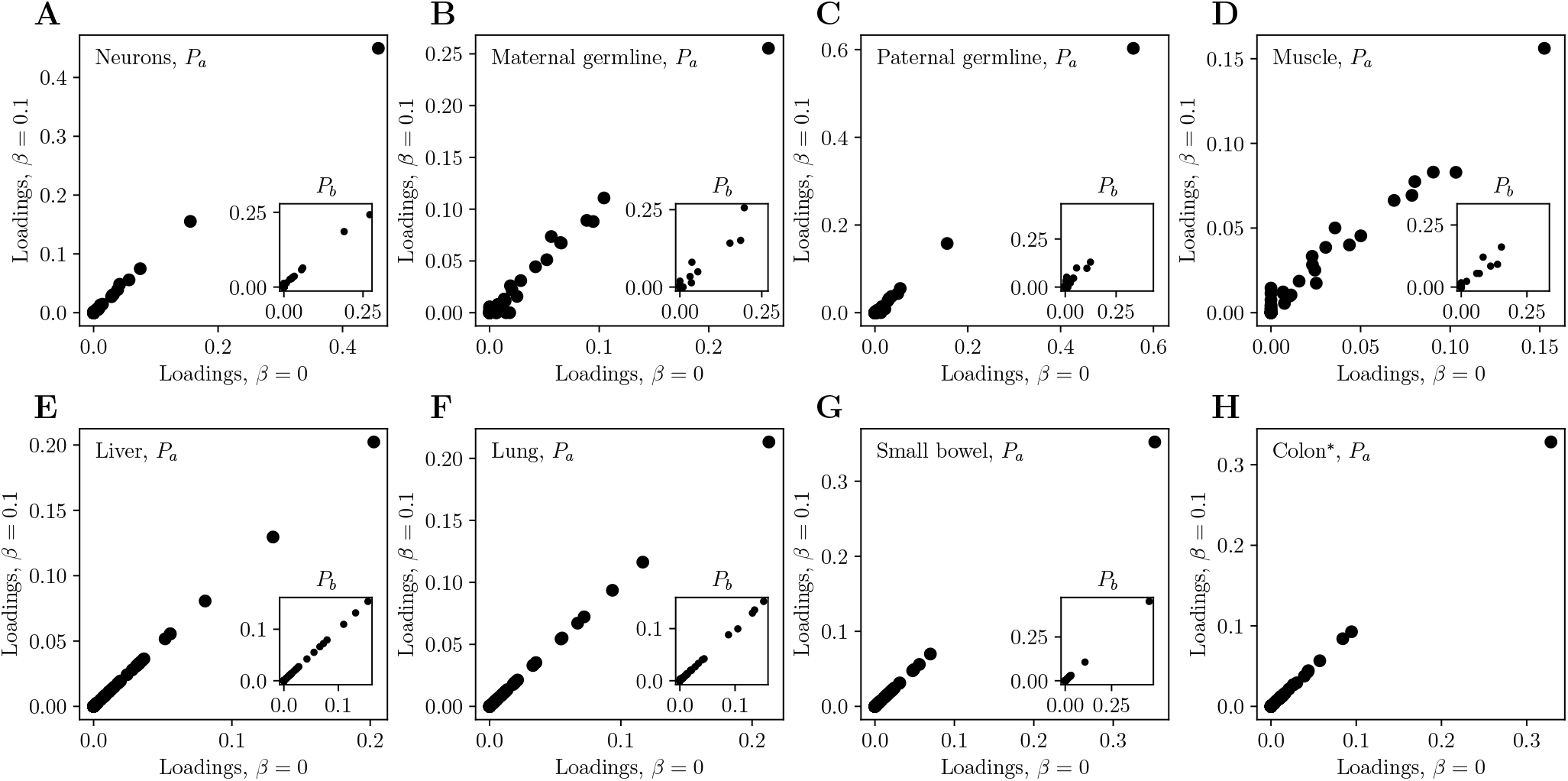
Inference results with weak regularization (*β* = 0.1) compared to no regularization (*β* = 0): loadings for the age-dependent component, *P*_*a*_, and the constant component, *P*_*b*_ (insets), in (A) neurons, (B) maternal germline mutations, (C) paternal germline mutations, (D) smooth muscle from bladder, (E) liver hepatocytes, (F) lung epithelium, (G) small bowel epithelium, (H) colon epithelium (^∗^for this dataset, in which the intercept is negative, we assume all signatures increase with age).

## Notes

### Competing Interest Statement

The authors have declared no competing interest.

## References

[1] Ségurel L, Wyman MJ, Przeworski M (2014) Determinants of mutation rate variation in the human germline. Annu. Rev. Genomics Hum. Genet. 15:47–70.

[2] Moore L, et al. (2021) The mutational landscape of human somatic and germline cells. Nature 597:381–386.

[3] Abascal F, et al. (2021) Somatic mutation landscapes at single-molecule resolution. Nature 593:405–410.

[4] Chatterjee N, Walker GC (2017) Mechanisms of DNA damage, repair, and mutagenesis. Environ. Mol. Mutagen. 58:235–263.

[5] Kucab JE, et al. (2019) A compendium of mutational signatures of environmental agents. Cell 177:821–836.e16.

[6] Nik-Zainal S, et al. (2012) Mutational processes molding the genomes of 21 breast cancers. Cell 149:979–993.

[7] Alexandrov LB, et al. (2013) Signatures of mutational processes in human cancer. Nature 500:415–421.

[8] Alexandrov LB, et al. (2020) The repertoire of mutational signatures in human cancer. Nature 578:94–101.

[9] Koh G, Degasperi A, Zou X, Momen S, Nik-Zainal S (2021) Mutational signatures: emerging concepts, caveats and clinical applications. Nat. Rev. Cancer 21:619–637.

[10] Tate JG, et al. (2019) COSMIC: the catalogue of somatic mutations in cancer. Nucleic Acids Res. 47:D941–D947.

[11] Cagan A, et al. (2022) Somatic mutation rates scale with lifespan across mammals. Nature 604:517–524.

[12] Rahbari R, et al. (2016) Timing, rates and spectra of human germline mutation. Nat. Genet. 48:126–133.

[13] Alexandrov LB, et al. (2015) Clock-like mutational processes in human somatic cells. Nat. Genet. 47:1402–1407.

[14] Matsuno Y, Kusumoto-Matsuo R, Manaka Y, Asai H, Yoshioka KI (2022) Echoed induction of nucleotide variants and chromosomal structural variants in cancer cells. Sci. Rep. 12:20964.

[15] Schiantarelli J, et al. (2022) Mutational footprint of platinum chemotherapy in a secondary thyroid cancer. JCO Precis Oncol 6:e2200183.

[16] Caballero M, Boos D, Koren A (2023) Cell-type specificity of the human mutation landscape with respect to DNA replication dynamics. Cell Genomics 3:100315.

[17] Guo Q, et al. (2022) The mutational signatures of formalin fixation on the human genome. Nat. Commun. 13:4487.

[18] Blokzijl F, et al. (2016) Tissue-specific mutation accumulation in human adult stem cells during life. Nature 538:260–264.

[19] Martínez-Jiménez F, et al. (2023) Pan-cancer whole-genome comparison of primary and metastatic solid tumours. Nature 618:333–341.

[20] Ivanov D, Hwang T, Sitko LK, Lee S, Gartner A (2023) Experimental systems for the analysis of mutational signatures: no ‘one-size-fits-all’ solution. Biochem. Soc. Trans. 51:1307–1317.

[21] Massaar S, Sanders MA (2023) The etiology of clonal mosaicism in human aging and disease. Aging Cancer 4:3–20.

[22] Yoshida K, et al. (2020) Tobacco smoking and somatic mutations in human bronchial epithelium. Nature 578:266–272.

[23] Ernst SM, et al. (2023) Tobacco Smoking-Related mutational signatures in classifying Smoking-Associated and Nonsmoking-Associated NSCLC. J. Thorac. Oncol. 18:487–498.

[24] Gao Z, Wyman MJ, Sella G, Przeworski M (2016) Interpreting the dependence of mutation rates on age and time. PLoS Biol. 14:e1002355.

[25] Tubbs A, Nussenzweig A (2017) Endogenous DNA damage as a source of genomic instability in cancer. Cell 168:644–656.

[26] Sanders MA, et al. (2021) Life without mismatch repair.

[27] Aitken SJ, et al. (2020) Pervasive lesion segregation shapes cancer genome evolution. Nature 583:265–270.

[28] Anderson CJ, et al. (2022) Strand-resolved mutagenicity of DNA damage and repair.

[29] Zhao L, Washington MT (2017) Translesion synthesis: Insights into the selection and switching of DNA polymerases. Genes 8.

[30] Powers KT, Washington MT (2018) Eukaryotic translesion synthesis: Choosing the right tool for the job. DNA Repair 71:127–134.

[31] Li GM (2008) Mechanisms and functions of DNA mismatch repair. Cell Res. 18:85–98.

[32] Ju YS, et al. (2017) Somatic mutations reveal asymmetric cellular dynamics in the early human embryo. Nature 543:714–718.

[33] Kaplanis J, et al. (2022) Genetic and chemotherapeutic influences on germline hypermutation. Nature 605:503–508.

[34] Brunner SF, et al. (2019) Somatic mutations and clonal dynamics in healthy and cirrhotic human liver. Nature 574:538–542.

[35] Wang Y, et al. (2023) APOBEC mutagenesis is a common process in normal human small intestine. Nat. Genet. 55:246–254.

[36] Halldorsson BV, et al. (2019) Characterizing mutagenic effects of recombination through a sequence-level genetic map. Science 363.

[37] Sasani TA, et al. (2019) Large, three-generation human families reveal post-zygotic mosaicism and variability in germline mutation accumulation. Elife 8.

[38] Lee-Six H, et al. (2019) The landscape of somatic mutation in normal colorectal epithelial cells. Nature 574:532–537.

[39] Alexandrov LB, et al. (2015) Clock-like mutational processes in human somatic cells. Nat. Genet. 47:1402–1407.

[40] Seplyarskiy VB, et al. (2021) Population sequencing data reveal a compendium of mutational processes in the human germ line. Science 373:1030–1035.

[41] Rodríguez-Nuevo A, et al. (2022) Oocytes maintain ROS-free mitochondrial metabolism by suppressing complex I. Nature 607:756–761.

[42] Alexandrov LB, et al. (2016) Mutational signatures associated with tobacco smoking in human cancer. Science 354:618–622.

[43] Miyaoka Y, et al. (2012) Hypertrophy and unconventional cell division of hepatocytes underlie liver regeneration. Curr. Biol. 22:1166–1175.

[44] Darwich AS, Aslam U, Ashcroft DM, Rostami-Hodjegan A (2014) Meta-analysis of the turnover of intestinal epithelia in preclinical animal species and humans. Drug Metab. Dispos. 42:2016–2022.

[45] Liu MH, et al. (2023) Single-strand mismatch and damage patterns revealed by single-molecule DNA sequencing.

[46] Zhang X, Mathews CK (1994) Effect of DNA cytosine methylation upon deamination-induced mutagenesis in a natural target sequence in duplex DNA. J. Biol. Chem. 269:7066–7069.

[47] Brown JA, Pack LR, Sanman LE, Suo Z (2011) Efficiency and fidelity of human DNA polymerases λ and β during gap-filling DNA synthesis. DNA Repair 10:24–33.

[48] Kunz C, Saito Y, Schär P (2009) DNA repair in mammalian cells: Mismatched repair: variations on a theme. Cell. Mol. Life Sci. 66:1021–1038.

[49] Hopfield JJ (1974) Kinetic proofreading: a new mechanism for reducing errors in biosynthetic processes requiring high specificity. Proc. Natl. Acad. Sci. U. S. A. 71:4135–4139.

[50] Bialek W (2012) Biophysics: Searching for Principles (Princeton University Press), Annotated edition edition.

[51] Gao Z, et al. (2019) Overlooked roles of DNA damage and maternal age in generating human germline mutations. Proc. Natl. Acad. Sci. U. S. A. 116:9491–9500.

[52] Moorjani P, Gao Z, Przeworski M (2016) Human germline mutation and the erratic evolutionary clock. PLoS Biol. 14:e2000744.

[53] Rey R, Josso N, Racine C (2020) Sexual Differentiation (MDText.com, Inc.).

[54] Grolleman JE, et al. (2019) Mutational signature analysis reveals NTHL1 deficiency to cause a multi-tumor phenotype. Cancer Cell 35:256–266.e5.

[55] Jónsson H, et al. (2017) Parental influence on human germline de novo mutations in 1,548 trios from iceland. Nature 549:519–522.

[56] Yan R, et al. (2021) Decoding dynamic epigenetic landscapes in human oocytes using single-cell multi-omics sequencing. Cell Stem Cell 28:1641–1656.e7.

[57] Gerstung M, et al. (2020) The evolutionary history of 2,658 cancers. Nature 578:122–128.

[58] Seplyarskiy VB, et al. (2019) Error-prone bypass of DNA lesions during lagging-strand replication is a common source of germline and cancer mutations. Nat. Genet. 51:36–41.

[59] Wu FL, et al. (2020) A comparison of humans and baboons suggests germline mutation rates do not track cell divisions. PLoS.

[60] Gelova SP, Doherty KN, Alasmar S, Chan K (2022) Intrinsic base substitution patterns in diverse species reveal links to cancer and metabolism. Genetics 222.

[61] Hwang DG, Green P (2004) Bayesian markov chain monte carlo sequence analysis reveals varying neutral substitution patterns in mammalian evolution. Proc. Natl. Acad. Sci. U. S. A. 101:13994–14001.

[62] Moorjani P, Amorim CEG, Arndt PF, Przeworski M (2016) Variation in the molecular clock of primates. Proc. Natl. Acad. Sci. U. S. A. 113:10607–10612.

[63] Hart RW, Setlow RB (1974) Correlation between deoxyribonucleic acid excision-repair and life-span in a number of mammalian species. Proc. Natl. Acad. Sci. U. S. A. 71:2169–2173.

[64] Tian X, et al. (2019) SIRT6 is responsible for more efficient DNA Double-Strand break repair in Long-Lived species. Cell 177:622–638.e22.

[65] Jahnke T, Huisinga W (2007) Solving the chemical master equation for monomolecular reaction systems analytically. J. Math. Biol. 54:1–26.

[66] Goldmann JM, et al. (2016) Parent-of-origin-specific signatures of de novo mutations. Nat. Genet. 48:935–939.

[67] Gehart H, Clevers H (2019) Tales from the crypt: new insights into intestinal stem cells. Nat. Rev. Gastroenterol. Hepatol. 16:19–34.

[68] Church DM, et al. (2011) Modernizing reference genome assemblies. PLoS Biol. 9:e1001091.

[69] Dempster AP, Laird NM, Rubin DB (1977) Maximum likelihood from incomplete data via theEMAlgorithm. J. R. Stat. Soc. 39:1–22.

[70] Figueiredo MAT, Jain AK (2002) Unsupervised learning of finite mixture models. IEEE Trans. Pattern Anal. Mach. Intell. 24:381–396.

[71] Virtanen P, et al. (2020) SciPy 1.0: fundamental algorithms for scientific computing in python. Nat. Methods 17:261–272.

[72] Schmutte C, Yang AS, Beart RW, Jones PA (1995) Base excision repair of U:G mismatches at a mutational hotspot in the p53 gene is more efficient than base excision repair of T:G mismatches in extracts of human colon tumors. Cancer Res. 55:3742–3746.

[73] Fang H, et al. (2021) Deficiency of replication-independent DNA mismatch repair drives a 5-methylcytosine deamination mutational signature in cancer. Sci Adv 7:eabg4398.

[74] Conti C, et al. (2007) Replication fork velocities at adjacent replication origins are coordinately modified during DNA replication in human cells. Mol. Biol. Cell 18:3059–3067.

[75] Tomkova M, McClellan M, Kriaucionis S, Schuster-Böckler B (2018) DNA replication and associated repair pathways are involved in the mutagenesis of methylated cytosine. DNA Repair 62:1–7.

[76] Seplyarskiy VB, Sunyaev S (2021) The origin of human mutation in light of genomic data. Nat. Rev. Genet. 22:672–686.

